# Insights on SARS-CoV-2’s Mutations for Evading Human Antibodies: Sacrifice and Survival

**DOI:** 10.1101/2021.02.06.430088

**Authors:** Binquan Luan, Tien Huynh

## Abstract

Recent mutations on the receptor binding domain (RBD) of the SARS-CoV-2’s spike protein have been manifested as the major cause of the wide and rapid spread of the virus. Especially, the variant B.1.351 in South Africa with the hallmark of triple mutations (N501Y, K417N and E484K) is worrisome. Quickly after the outbreak of this new variant, several studies showed that both N501Y and E484K can enhance the binding between RBD and the human ACE2 receptor. However, the mutation K417N seems to be unfavorable because it removes one interfacial salt-bridge. So far, it is still not well understood why the K417N mutation is selected in the viral evolution. Here, we show that despite the loss in the binding affinity (1.48 kcal/mol) between RBD and ACE2 the K417N mutation abolishes a buried interfacial salt-bridge between RBD and the neutralizing antibody CB6 and thus substantially reduces their binding energy by 9.59 kcal/mol, facilitating the variants to efficiently elude CB6 (as well as many other antibodies). Thus, when proliferating from person to person the virus might have adapted to the human immune system through evasive mutations. Taking into account limited and relevant experimental works in the field, we show that our theoretical predictions are consistent with existing experimental findings. By harnessing the revealed molecular mechanism for variants, it becomes feasible to redesign therapeutic antibodies accordingly to make them more efficacious.

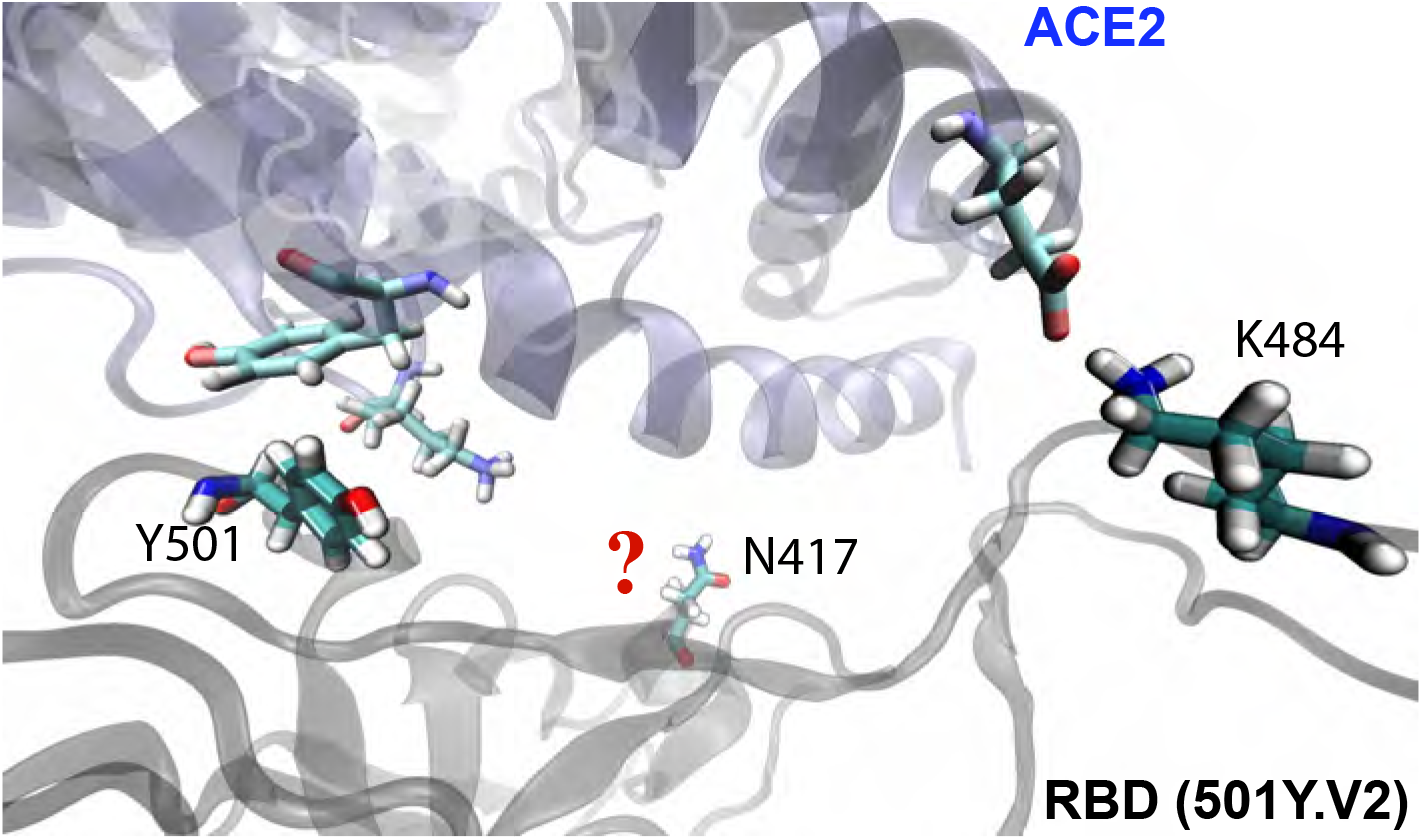

---

Severe acute respiratory syndrome coronavirus 2 (SARS-CoV-2)^1^ is a positive-sense single-stranded RNA virus that causes the ongoing coronavirus disease 2019 (COVID-19) pandemic. Because SARS-CoV-2 is a highly transmissible and pathogenic novel coronavirus, the global morbidity and mortality rates of COVID-19 have been quite substantial since it was first detected at the end of 2019.^2^ This has brought together researchers from the international scientific community to combat COVID-19 with extraordinary efforts. Although we have come a long way in a short period of time, it is still challenging to have the pandemic under full control especially with the recent discovery of several new variants of SARS-CoV-2, such as B.1.1.7 (or 501Y.V1) from U.K., B.1.351 (or 501Y.V2) from South Africa and P.1 from Brazil. With the increased transmissibility and hence rapid spreading of these newly identified lineages, serious concerns have been raised over whether they could undermine the currently available vaccines or natural immunity of people previously infected with COVID-19.^3^

Experimental studies have demonstrated that the angiotensin converting enzyme 2 (ACE2), a protein expressed on the surface of human cells in various organs, plays a crucial role in the viral infection of SARS-CoV-2.^4–6^ As a prelude of SARS-CoV-2’s cell entry, the receptor binding domain (RBD) of the spike glycoprotein (S-protein)^7^ on the virion surface binds ACE2 on a host cell. Therefore, the S-protein of SARS-CoV-2 becomes a primary target of the host immune system and is used as the leading antigen for vaccine development. Despite being detected from three different continents, the fact that the three aforementioned SARS-CoV-2 variants share some of the same mutations at the S-protein’s RBD suggests that these mutations might have conferred an evolutionary advantage to the virus. While various mutations as well as deletions outside the RBD of S-protein are also important for the enhanced fitness (of variants) for entering host cells, here we focus on mutations in RBD that harbors the binding site of ACE2 and the epitopes for neutralizing monoclonal antibodies (mAbs).

For the U.K. variant (with the N501Y mutation on RBD), experimental studies have demonstrated that the reproductive number which measures its infectiousness is about 0.4 to 0.7 higher than other strains of the virus^8^ and determined recently it is unlikely to escape the BNT162b2-vaccine–mediated protection.^9^ Our previous work^10^ showed that this mutation in the U.K. variant can increase the RBD’s binding affinity with ACE2, but has no obvious effect on mAbs. Besides the N501Y mutation, the South Africa and Brazil variants also contain the K417N and E484K mutations. Recent studies showed that the E484K mutation can not only enhance the RBD-ACE2 binding^11^ but also help virus escape the therapeutically relevant mAbs.^12^ However, reflected by the paucity of existing work the significance of the K417N mutation so far is still elusive, which prevents us from fully understanding these new variants’ infection mechanism. As shown in Fig. 1a, in the wild-type RBD, K417 forms a salt-bridge with D30 in ACE2. Thus, the K417N mutation resulting in the abolishment of this favorable interfacial interaction (i.e. reducing the RBD-ACE2 binding affinity as also being verified in experiment^11^) is highly unintuitive. Here, we are motivated to investigate the molecular mechanism of the K417N mutation, in order to unveil its key benefits for the virus to evolve through this path.

**Figure 1:**
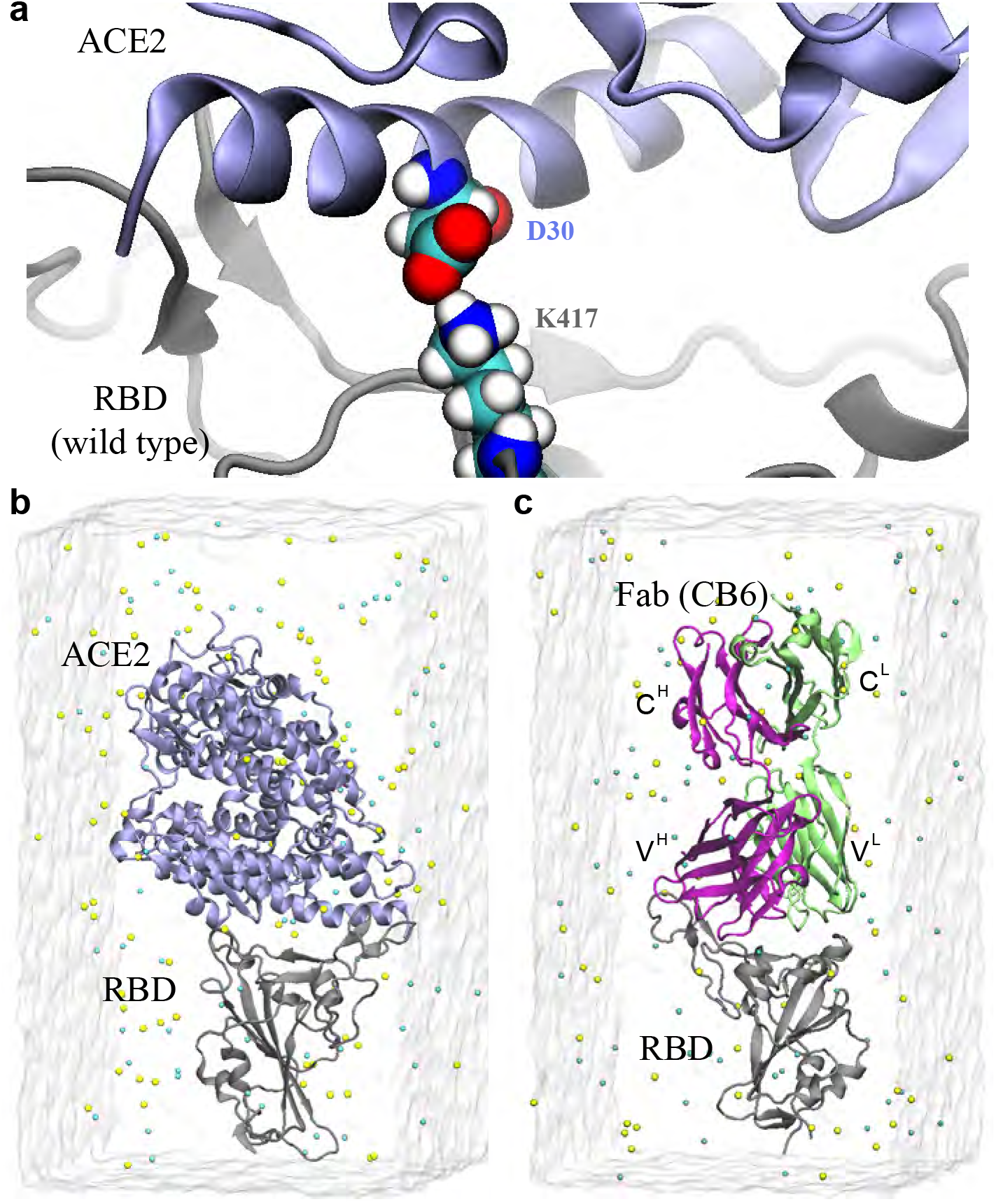
MD simulation systems. a) Illustration of the salt-bridge formed by D30 in ACE and K417 in RBD at the interface. b) Simulation setup for the RBD-ACE2 complex. c) Simulation setup for the RBD-CB6 complex. Proteins are in the cartoon representation, with RBD in gray, ACE2 in blue, the heavy (V^*H*^ and C^*H*^) and light (V^*L*^ and C^*L*^) chains of the CB6’s Fab in purple and green respectively. Na^+^ and Cl^−^ are shown as yellow and cyan balls respectively. Water is shown transparently.

Complementary to experimental efforts, the all-atom molecular dynamics (MD) simulations with sophisticated and well calibrated force fields have been widely used to image nanoscale events and investigate the molecular mechanism of proteins.^13–16^ In this work, we conducted a computational analysis on the K417N mutation in the South Africa variant using MD simulations with explicit solvent, aiming to gain a better understanding of its underlying molecular mechanism. Besides the RBD-ACE2 interaction, we also investigated the RBD’s interaction with mAbs. In particularly, we explored the mAb CB6 that recognizes an epitope site in the RBD overlapping the binding site of ACE2, and investigated its binding competition with ACE2 over RBD. Our results may help provide invaluable insights on why K417N has been selected in the viral evolution and inspire a better design of more efficacious mAbs for treating COVID-19 patients infected with the new SARS-CoV-2 variants.

Figure 1b illustrates the simulation system for modeling the interaction between ACE2 and RBD (see the Methods section for detailed simulation protocols). Briefly, taken from the crystallographic structure (PDB: 6M0J), the RBD-ACE2 complex’s structure was solvated in a 0.15 M NaCl electrolyte. Similarly, as shown in Fig. 1c, we built the simulation system for the complex of RBD and one Fab of the human antibody CB6 with their atomic coordinates taken from the crystallographic structure (PDB: 7C01). Hereafter, we simply refer the Fab as CB6.

During the 350 ns or so MD simulation, the RBD-ACE2 complex starting from the structure in the crystal environment was equilibrated in the physiology-like environment (a 0.15 M electrolyte). Figure 2a shows the root-mean-square-deviation (RMSD) of protein-backbone atoms in RBD bound either with ACE2 or CB6. After about 40 ns, the RMSD values saturated at around 1.7 Å for RBD bound on CB6 and 1.5 Å for RBD bound on ACE2. These small RMSD values suggest that the secondary structure of modeled RBD (with four internal disulfide bonds) were very stable and properly equilibrated. We also calculated the interfacial contact areas for the complexes using the solvent accessible surface area method.^17^ On average, the contact area is about 8.6 nm^2^ between RBD and ACE2 while the one for the RBD-CB6 contact is larger and is about 10.3 nm^2^. During the entire simulation time, values of contact areas fluctuated around the mean, indicating that these two complex-structures were stable and equilibrated in the electrolyte.

**Figure 2:**
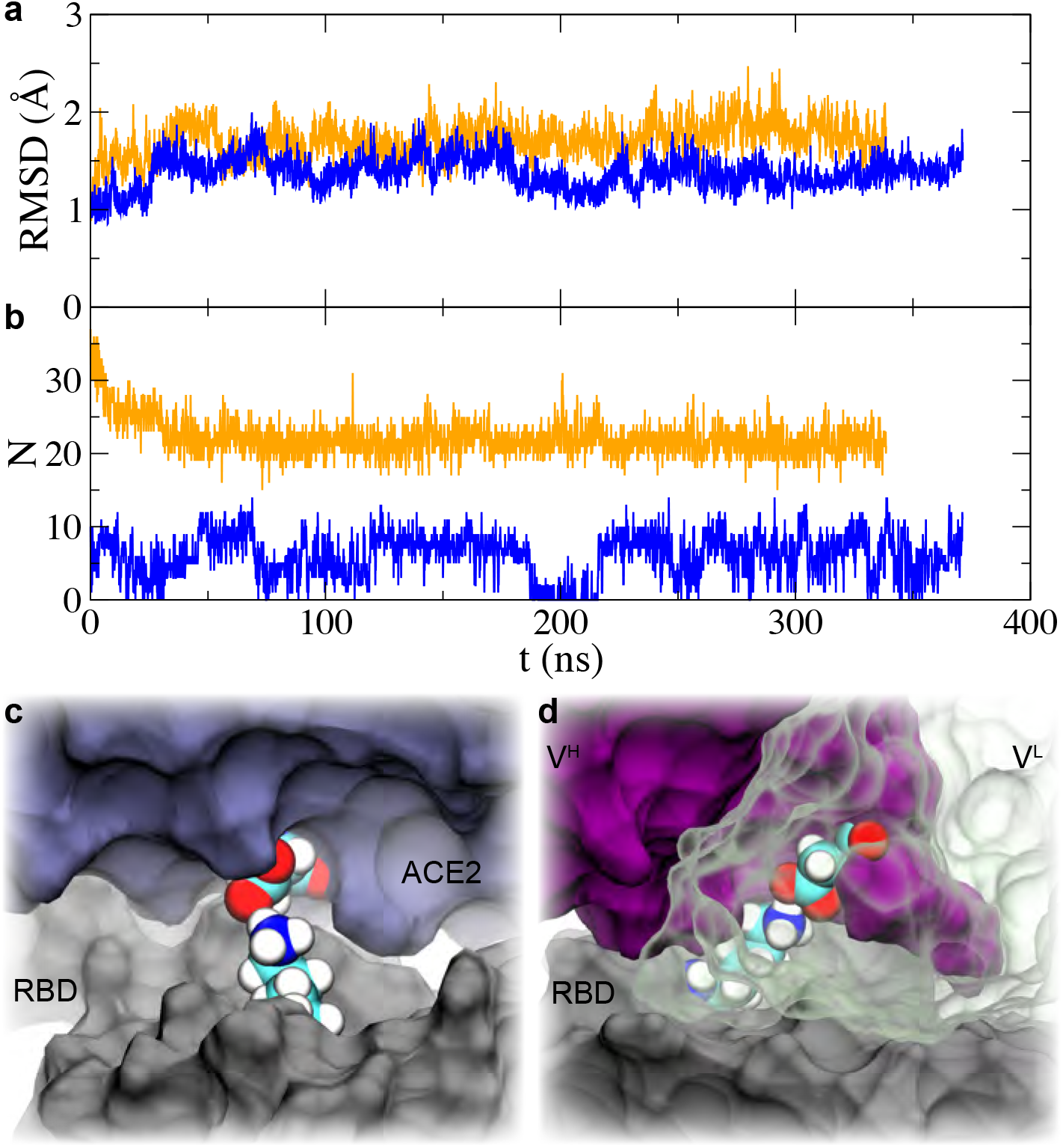
MD simulations of both RBD-ACE2 and RBD-CB6 complexes. a) Time-dependent RMSDs of RBDs in simulated complexes, RBD-ACE2 (blue) and RBD-CB6 (orange). b) Time-dependent numbers of heavy (or non-hydrogen) atoms in ACE2 (blue) or CB6 (orange) that are within 5 Å of K417 in RBD. c) Illustration of the salt-bridge formed by K417 in RBD and D30 in ACE2 that is present on the complex’s surface (i.e. being exposed to water). d) Illustration of the salt-bridge formed by K417 in RBD and D104 in CB6 (V^*H*^) that is buried inside the complex. RBD, ACE2, V^*H*^ and V^*L*^ of CB6 are in the representation of molecular surface. V^*L*^ of CB6 is transparent to allow viewing of the buried salt-bridge.

By examining the simulation trajectories, we found that K417 indeed played an important role in stabilizing the RBD-ACE2 and RBD-CB6 complexes. In Fig. 2b, we highlight the time-dependent number *N* of heavy (or non-hydrogen) atoms in ACE2 or CB6 that were within 5 Å of K417 in RBD. Notably, K417 in RBD interacts with many more atoms in CB6 than in ACE2. The average saturated number N for the RBD-CB6 complex is about 23 atoms (orange line in Fig. 2b), while the one for the RBD-ACE2 complex is only about 6 atoms (blue line in Fig. 2b). Additionally, the number N for the RBD-ACE2 complex fluctuates much more than the one for the RBD-CB6 complex, which indicates that the interaction in the latter complex is more stable.

Among those residues (in ACE2 or CB6) in contact with K417 in RBD, we discovered two key salt-bridges. One is formed by K417 in RBD and D30 in ACE2 (as shown in Fig. 1a). We illustrate in Fig. 2c that this salt-bridge is on the surface of the RBD-ACE2 complex. Thus, this salt-bridge is exposed to water and constantly disrupted by polar water molecules, accounting for the observed fluctuations in N (blue line in Fig. 2b). The other one is formed by K417 in RBD and D104 in CB6 as shown in Fig. 2d. Remarkably, this salt-bridge is buried among the heterotrimer composed of RBD and two variable domains of the heavy chain *V^H^* and light chain *V^L^* of CB6. This salt-bridge buried inside the protein complex was very stable during the simulation, consistent with the nearly constant number *N* in Fig. 2b (orange line).

To further demonstrate the stability of these two salt-bridges, we define the distance *D* between the NZ atom in the lysine (K) and the CG atom in the aspartate (D), as shown in the inset of Fig. 3a. The histograms in Fig. 3a show the probability distributions of the distance *D*. For the salt-bridge in RBD-CB6, the most probable distance in the single-peak distribution is about 3.2 Å, corroborating that this buried salt-bridge is very stable. For the salt-bridge in RBD-ACE2, the most probable distance is in the first peak and is about 3.5 Å. This larger distance (as compared with the one for the buried salt-bridge) reflects the reduced electrostatic interaction inside the water-exposed salt-bridge. Furthermore, an additional peak at larger distances (~5.5 Å) is an indication of a state of the broken saltbridge. Indeed, we observed in the simulation trajectory that this salt-bridge is susceptible to polar water molecules and can break and reform from time to time.

**Figure 3:**
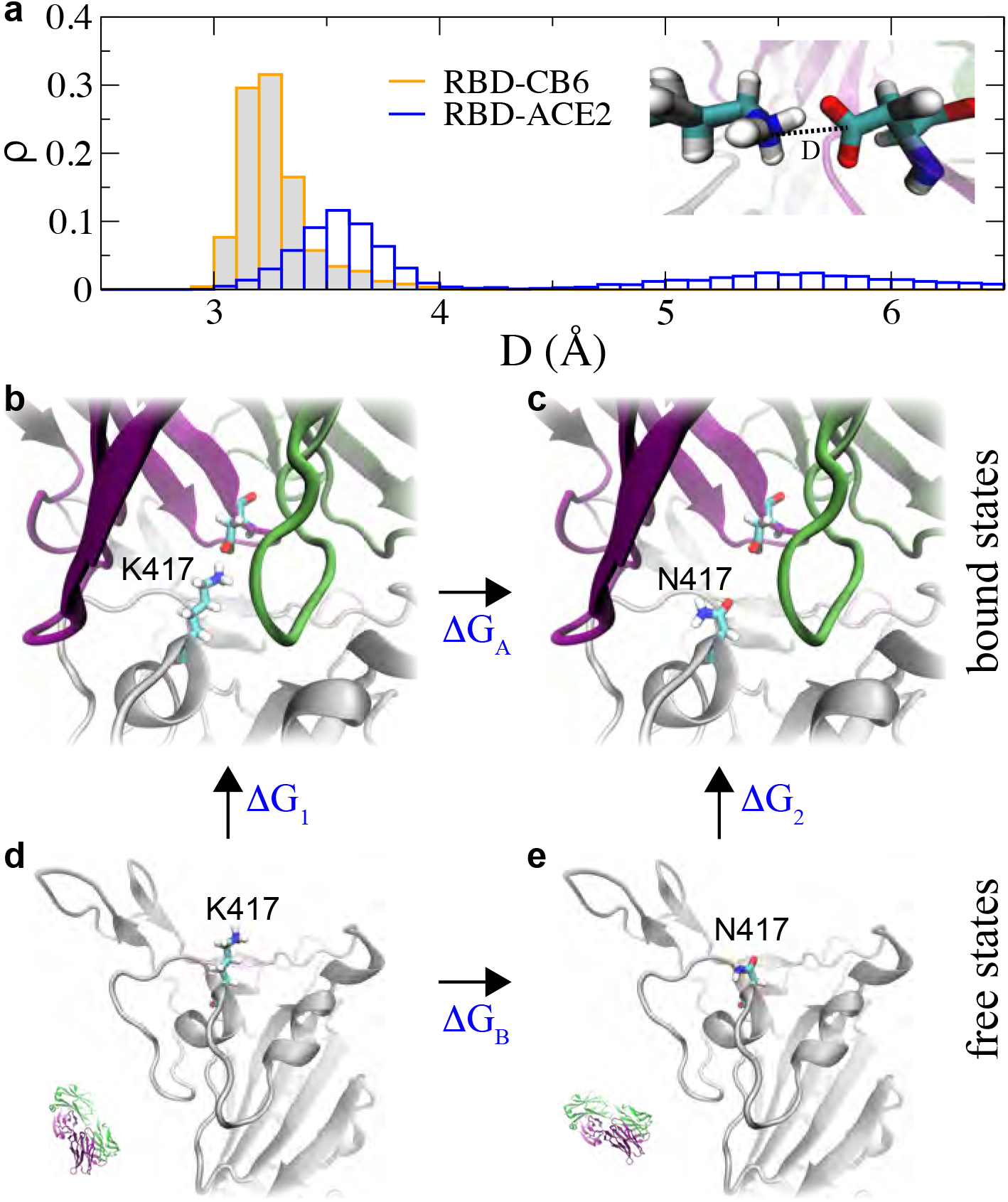
Characterizing salt-bridges at the RBD-CB6 and RBD-ACE2 interfaces. a) Distribution of the distance *D* between a pair of residues in a salt-bridge, at the RBD-CB6 interface (orange) and at the RBD-ACE2 interface (blue). The *inset* shows the definition of the distance *D*. b-e) The thermal dynamic cycle to obtain the binding free energy change induced by the K417N mutation. The mutation in the bound and free states are shown in (b-c) and (d-e) respectively.

Therefore, in the wild-type RBD, K417 interacts more strongly with D104 in CB6 than with D30 in ACE2. The calculations of the binding free energy change induced by the K417N mutation were accomplished using the free energy perturbation (FEP) method.^18^ As required in FEP calculations, We performed 177-ns-long MD simulations of RBD in a 0.15 M NaCl electrolyte (a free state), as shown in Fig. S1 in Supporting Information. With protein structures for both bound (or complex) and free states in respective MD simulations, we employed the FEP alchemy method to calculate the binding free energy difference for the K417N mutation on the RBD, using the thermodynamic cycle shown in Figs. 3c-3e. By definition, binding free energy changes for RBD with either ACE2 or CB6 (due to the K417N mutation) can be obtained as ΔΔ*G* = Δ*G*_2_ – Δ*G*_1_, (see Fig. 3). In practice, it is not easy to directly calculate Δ*G*_1_ and Δ*G*_2_, which can be circumvented by computing Δ*G_A_* and Δ*G_B_* instead using the thermodynamic cycle (see Fig. 3). Therefore, ΔΔ*G* = Δ*G_A_* – Δ*G_B_*. Through the ensemble average,^18^ Δ*G_A_* and Δ*G_B_* can be calculated theoretically (see Method section) as 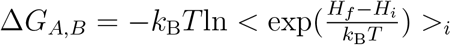, where *k_B_* is the Boltzmann constant; *T* the temperature; *H_i_* and *H_f_* the Hamiltonians for the initial (*i*) stage with K417 (see Figs. 3b and 3d) and the final (*f*) stage with N417 (see Figs. 3c and 3e), respectively.

Results from the FEP calculations are summarized in Tab. 1. In the free state (Figs. 3d and 3e), the K417N mutation yielded a free-energy change Δ*G_B_* of −32.08 kcal/mol. In the bound state for the RBD-CB6 (Figs. 3b and 3c), the free energy change Δ*G_A_*=−22.49 kcal/mol. Taking all together, the obtained value of ΔΔ*G* is 9.59 kcal/mol, suggesting that the K417N mutation significantly reduced the binding affinity between RBD and CB6. Similarly, for the RBD-ACE2 complex, we have Δ*G_A_*=−30.60 kcal/mol and consequently ΔΔ*G* is 1.48 kcal/mol. Thus, the K417N mutation reduced the binding affinity between RBD and ACE2 as well, but the reduction is about 6.5 time less than that between RBD and CB6. Previously, it was demonstrated in experiment that the K417N mutation weakened the binding affinity between RBD and ACE2,^11^ which is consistent with our simulation results. The noticeably small errors listed in Tab. 1 manifest the accuracy and convergence afforded by the FEP methodology.

**Table 1:**
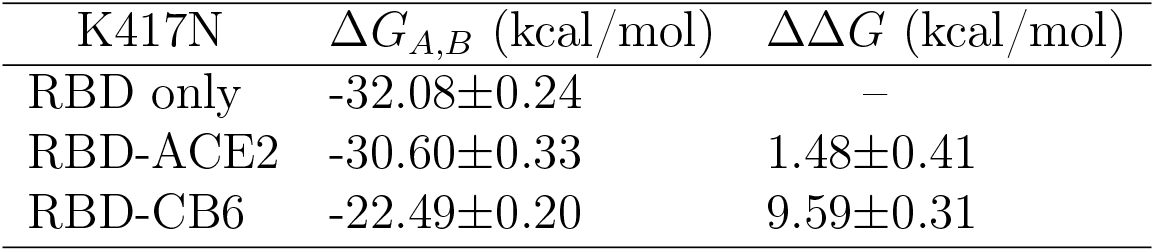
Values of ΔΔ*G* for the K417N mutation, in RBD’s binding with ACE2 and CB6 (mAb).

| K417N | $\Delta G_{A,B}$ (kcal/mol) | $\Delta\Delta G$ (kcal/mol) |
| --- | --- | --- |
| RBD only | -32.08±0.24 | – |
| RBD-ACE2 | -30.60±0.33 | 1.48±0.41 |
| RBD-CB6 | -22.49±0.20 | 9.59±0.31 |

The free energy loss due to the K417N mutation is mainly caused by the change in electrostatic energy. The dielectric constant *ϵ*_*p*_ inside a protein is approximately 4,^19^ while the dielectric constant *ϵ*_*w*_ of water is about 90 for the TIP3P model.^20^ If we approximate the protein-water interface with a planar surface separating two dielectric materials (*ϵ*_*p*_ and *ϵ*_*w*_), the effective dielectric constant on the interface can be calculated as (*ϵ*_*p*_+ϵ_*w*_)/2, from a simple continuum electrostatics calculation. Thus, the ratio *γ* of electrostatic energy changes for salt-bridges on the protein surface and in water can be roughly estimated as (ϵ_*p*_+ϵ_*w*_)/2ϵ_*w*_~11.8 (larger than our simulation result γ=6.5). However, the more realistic explicit model with atomic details^21^ predicts that the average dielectric constant on the protein-water interface is about 20-30, yielding that γ ~5-7.5, which is in an excellent agreement with our numerical result.

More importantly, the phenomenon of weakening the binding affinity between RBD-CB6 observed in the K417N mutation is not unique. After comparing some crystal structures for the complex of RBD (of the SARS-CoV-2’s spike protein) and human antibodies, surprisingly we noticed that K417 in RBD can form a buried salt-bridge with either a glutamate or an aspartate in four other human antibodies as highlighted in IGHV-53 (Fig. 4a), C1AB3 (Fig. 4b), BD-236 (Fig. 4c) and CC12.1 (Fig. 4d). Thus, it is likely that all these antibodies may not be able to neutralize the 501Y.V2 variant. It is worth noting that while writing this paper, an experimental preprint posted on bioRxiv demonstrating that the South Africa variant can evade the human antibodies CB6 (Fig. 1c, also known as LyCoV016) and CC12.1 (Fig. 4d) that are known to neutralize the wild-type SARS-CoV-2.^12^ This experimental result further confirms our predictions. Because K417 in RBD can form a buried salt-bridge with a glutamate/aspartate in a class of human antibodies, we speculate that K417 is the Achilles’ heel of the virus and can be easily targeted by many human antibodies. Therefore, the K417N mutation could be the evidence of viral adaptation to the human immune system, or the natural selection under the “pressure” of human neutralizing mAbs.

**Figure 4:**
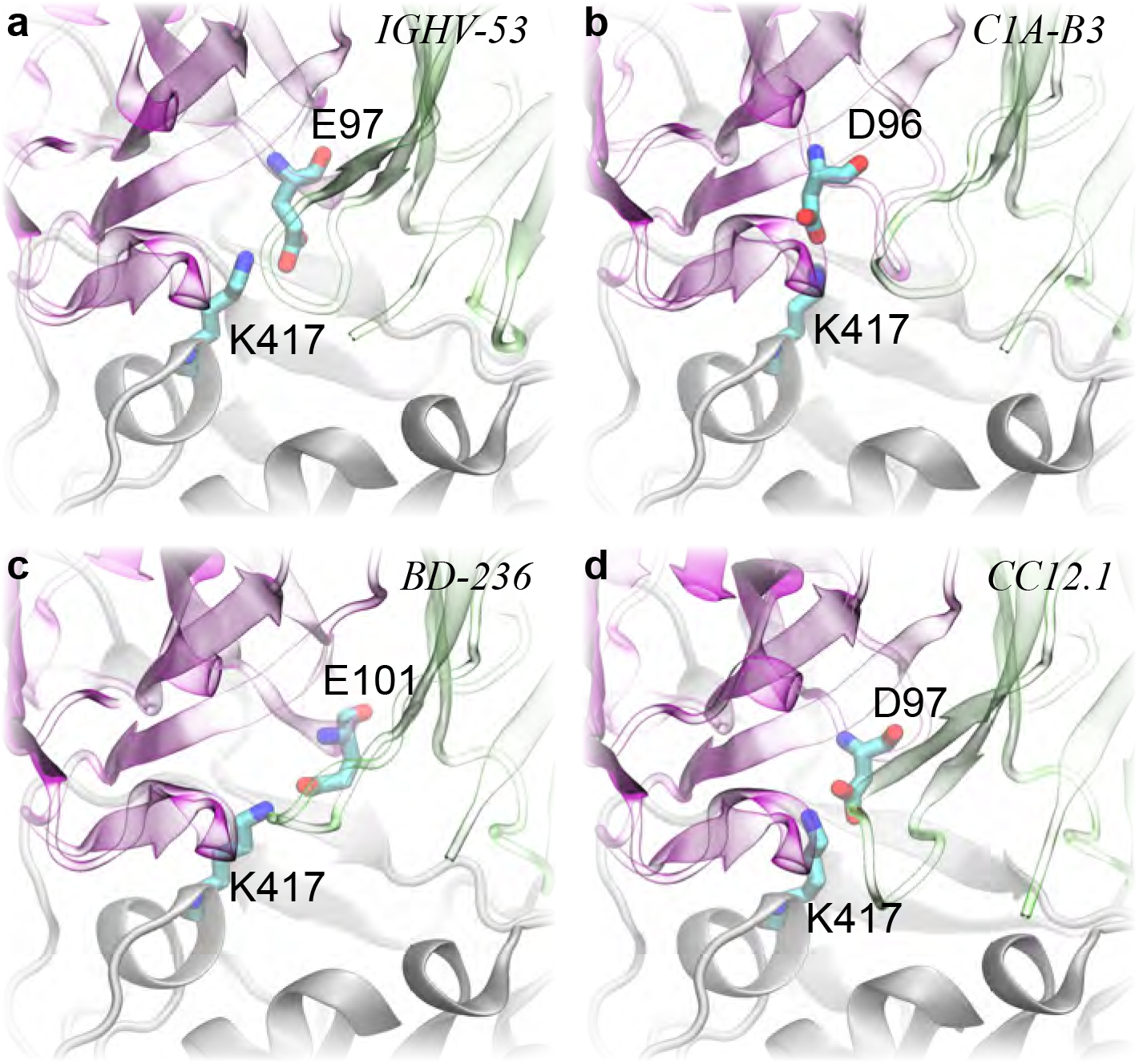
Illustrations of various salt-bridges buried at the RBD-antibody interfaces. a) The neutralizing antibody IGHV-53 (PDB: 7JMO) for SARS-CoV-2’s RBD; b) The neutralizing antibody C1A-B3 (PDB: 7B3O) for SARS-CoV-2’s RBD; c) The neutralizing antibody BD-236 (PDB: 7CHB) for SARS-CoV-2’s RBD; d) The neutralizing antibody CC12.1 (PDB: 6XC2) for SARS-CoV-2’s RBD. The RBD, V^*L*^ and V^*H*^ of various antibodies are colored in gray, green and purple respectively. Residues forming a salt-bridge at the RBD-antibody interface are in the stick representation. V^*L*^ and V^*H*^ of various antibodies are shown transparently to allow viewing of buried salt-bridges.

Although it is counterintuitive to learn that the K417N mutation actually lowering the RBD’s binding affinity with ACE2 by 1.48 kcal/mol, it is in fact beneficial for RBD to escape the human antibody CB6 by dramatically reducing the binding affinity by 9.59 kcal/mol. Because of the loss in the binding affinity, the variant with this single mutation are less competitive than the wild-type SARS-CoV-2 virus, in terms of the ability for entering host cells. So far there is no evidence that this particular mutation occurs alone in any known variants of SARS-CoV-2. In the South Africa variant (501Y.V2), actually there are two more mutations, N501Y and E484K. To explore their contributions, we carried out MD simulation for the RBD-ACE2 complex with all three mutations (K417N, N501Y and E484K), as shown in Fig. S2a in Supporting Information. After about 340-ns MD simulation, both N501Y and E484K are found to improve the interfacial binding between RBD and ACE2.

The N501Y mutation also occurred in the U.K. variant (501Y.V1) and our previous modeling of this variant showed that this mutation can increase the binding affinity by forming hydrophobic interfacial interactions.^10^ For our current simulation with the triple mutations (Fig. S2a), we observed the same improved binding structure (see Fig. S2b and Movie S1 in the Supporting Information). For the E484K mutation, after about 190ns simulation, K484 moved toward ACE2 to form a salt-bridge exposed on the complex’s surface with E75 (see Figs. S2b and S2c, and Movie S1 in the Supporting Information). Therefore, the gain from the improved RBD-ACE2 binding resulted from both N501Y and E484K mutations might be enough to compensate the loss caused by the K417N mutation, and is likely to yield an even stronger interaction with ACE2 than the wild-type virus.

In summary, we have shown that K417 plays an important role in the binding between RBD and ACE2/CB6, by forming interfacial salt-bridges. The salt-bridge between K417 in RBD and D104 in CB6 is buried inside the complex and is therefore much more stable than the water exposed one formed between K417 in RBD and D30 in ACE2. Thus, the K417N mutation can weaken the RBD’s binding with CB6 much more than with ACE2. More dramatically, the K417N mutation allows the variant to escape from many human antibodies other than CB6 by removing a salt-bridge buried in the RBD-antibody interface. Interestingly, the virus with the K417N mutation seems to scarify its binding affinity with ACE2 in order to survive the antibodies’ attack. This strategy of eluding human antibodies might have been adopted by SARS-CoV as well, because after the sequence alignment the residue in SARS-CoV corresponding to K417 in SARS-CoV-2 is the valine (V).

On the other hand, a recent study has demonstrated that immune systems of human may also quickly respond to these new variants and produce corresponding neutralizing antibodies that contain more somatic hypermutation, increased potency and resistance to RBD mutations.^22^ How the continued evolution of the humoral response to recent SARS-CoV-2 variants warrants further studies. Additionally, with the revealed mechanism underlying the K417N mutation, it is possible to design more efficacious antibody cocktails to treat COVID-19 patients infected with the variant 501Y.V2 as well as the recently discovered variant P.1 in Brazil.

## Methods

### MD simulations

We carried out all-atom MD simulations for both the bound (the RBD-ACE2 complex) and free (stand alone RBD) states using the NAMD2.13 package^23^ running on the IBM Power Cluster. To model the RBD-ACE2 and RBD-CB6 complexes, we first obtained the crystal structures with PDB codes 6M0J^24^ and 7C01^25^ from the protein data bank respectively, and then solvated them in a rectangular water box that measures about 80×80×135 Å^3^. A bound Zn^2+^ ions were included in the RBD-ACE2 complex. Na^+^ and Cl^-^ were added to neutralize the entire simulation system and set the ion concentration to be 0.15 M (Figs. 1b and 1c). Each built system was first minimized for 10 ps and further equilibrated for 1000 ps in the NPT ensemble (*P* ~ 1 bar and *T* ~ 300 K), with atoms in the backbones harmonically constrained (spring constant *k*=1 kcal/mol/Å^2^). The production run was performed in the NVT ensemble, only atoms in the backbones of ACE2 that are far away from the RBD (residues 110 to 290, 430 to 510 and 580 to 615 in ACE2) were constrained, preventing the whole complex from rotating out of the water box. The similar approach was applied in the production run for the RBD-CB6 complex. We also performed MD simulation for the RBD alone in the 0.15 M NaCl electrolyte (a free state) using the same protocol (Fig. S1 in Supporting Information).

After equilibrating structures in bound and free states, we carried out free energy perturbation (FEP) calculations.^18^ In the perturbation method, many intermediate stages (denoted by λ) whose Hamiltonian *H*(λ)=λ*H_f_*+(1-λ)*H_i_* are inserted between initial and final states to yield a high accuracy. With the softcore potential enabled, λ in each FEP calculation for Δ*G_A_* or Δ*G_B_* varies from 0 to 1.0 in 20 perturbation windows (lasting 300 ps in each window), yielding gradual (and progressive) annihilation and exnihilation processes for K417 and N417, respectively. We followed our previous FEP protocol^26^ to obtain the means and errors for Δ*G_A_* and Δ*G_B_*.

We applied the CHARMM36m force field^27^ for proteins, the TIP3P model^28,29^ for water, the standard force field^30^ for Na^+^, Zn^2+^ and Cl^−^. The periodic boundary conditions (PBC) were used in all three dimensions. Long-range Coulomb interactions were calculated using particle-mesh Ewald (PME) full electrostatics with the grid size about 1 Å in each dimension. The pair-wise van der Waals (vdW) energies were calculated using a smooth (10-12 Å) cutoff. The temperature *T* was maintained at 300 K by applying the Langevin thermostat,^31^ while the pressure was kept constant at 1 bar using the Nosé-Hoover method.^32^ With the SETTLE algorithm^33^ enabled to keep all bonds rigid, the simulation time-step was set to be 2 fs for bonded and non-bonded (including vdW, angle, improper and dihedral) interactions, and electric interactions were calculated every 4 fs, with the multiple time-step algorithm.^34^

## Acknowledgement

T.H and B.L. gratefully acknowledge the computing resource from the IBM Cognitive Computing Program and useful discussion with Haoran Wang.

## Supporting Information Available

MD simulation for RBD-only (a free state); MD simulation for the complex comprising human ACE2 and RBD of S-protein from the South Africa variant; Movie (SA.mov) showing dynamic interfacial interaction between human ACE2 and RBD of S-protein from the South Africa variant.

